# Grazing results in mobilization of spherulous cells and re-allocation of secondary metabolites to the surface in the sponge *Aplysina aerophoba*

**DOI:** 10.1101/2020.11.20.391169

**Authors:** Yu-Chen Wu, María García-Altares, Berta Pintó, Marta Ribes, Ute Hentschel, Lucía Pita

## Abstract

On the sea floor, prey and predator commonly engage in a chemical warfare. Here, sponges thrive due to their specific and diverse chemical arsenal. Yet, some animals use these chemically-defended organisms as food and home. Most research on sponge chemical ecology has characterized crude extracts and investigated defences against generalist predators like fish. Consequently, we know little about intraindividual chemical dynamics and responses to specialist grazers. Here, we studied the response of the sponge *Aplysina aerophoba* to grazing by the opistobranch *Tylodina perversa*, in comparison to mechanical damage, at the cellular (via microscopy) and chemical level (via matrix-assisted laser desorption/ionization imaging mass spectrometry). We characterized the distribution of two major brominated compounds in *A. aerophoba*, aerophobin-2 and aeroplysinin-1, and identified a generalized wounding response that was similar in both wounding treatments: (i) brominated compound-carrying cells (spherulous cells) accumulated at the wound and (ii) secondary metabolites reallocated to the sponge surface. Upon mechanical damage, the wound turned dark due to oxidized compounds, causing *T. perversa* deterrence. During grazing, *T. perversa’s* way of feeding prevented oxidation. Thus, the sponge has not evolved a specific response to this specialist predator, but rather relies on rapid regeneration and flexible allocation of constitutive defences.

## INTRODUCTION

Many sessile marine organisms have developed chemical defences to avoid predators, compete for space, and prevent fouling and colonization by pathogens and opportunistic microbes. These defences may be constitutive (always present in the organisms), inducible upon stimuli (synthesized anew), or activated when the organism is attacked (converted from a non- or less toxic precursor to a potent toxin) ^1–3^. Nevertheless, some specialized predators have overcome these chemical defences in order to take advantage of an otherwise unexploited food source. These specialists are often small animals (mesograzers) for which their prey also becomes their habitat since, despite its poor nutritional value, feeding on chemically-defended organisms provides the additional benefit of protection ^1,4,5^. Thus, chemical defences mediate species-species interactions and determine organisms’ success, shaping the diversity and function of benthic communities ^6^.

Sponges represent a prominent example of chemically-defended marine organisms with great ecological success. They have been extensively studied because of the potential medical and biotechnological application of their secondary metabolites and constitute the richest source of marine natural products ^7–10^. Each sponge species produces a specific, yet diverse chemical arsenal with fish-deterrent, antifouling and antimicrobial properties, to name a few ^3,4,11–13^. However, a great variety of beautiful opistobranchs are specialized in living and feeding on one or a narrow range of sponge species. These specialized sea slugs defend themselves by accumulating and modifying the secondary metabolites they acquire from the sponges they eat ^4^. Despite these well-known associations, and the extensive literature on sponge chemical defences, we know little about the response of sponges to grazing by these specialists and whether it differs from the response to predation by generalists or wounding. Moreover, most research has focused on the concentration and bioactivity of secondary metabolites in crude extracts and we lack spatial resolution on the distribution of compounds within individual sponges.

Here, we investigated the interaction between the sponge *Aplysina aerophoba* (Nardo, 1833) and the sea slug *Tylodina perversa* (Gmelin, 1791). Sponges of the genus *Aplysina* serve as models for sponge chemical ecology ^14–19^ and have stimulated a heated debate on activated defences in sponges. *Aplysina* sponges are widely distributed and present similar chemical profiles ^20,21^, dominated by several brominated isoxazoline alkaloids. In *A. aerophoba*, the most abundant ones are aerophobin-2, isofistularin-3, and aplysinamisin-1, and showed repellence activity against marine fishes ^22^. These compounds are traditionally considered precursors of the smaller, more cytotoxic compounds aeroplysinin-1 and its structurally related dienone ^13,23–25^ and it was proposed that this is an enzymatically-mediated transformation ^23,24^. These bioconversion products display antimicrobial activity upon wounding or stress ^19,25^. However, those studies suggesting induced enzymatic transformation have been highly criticised, mainly because of possible methodological artefacts and because experiments on two Caribbean *Aplysina* species did not support activated antipredatory defences ^26^. Yet, a nitrile hydratase that specifically takes aeroplysinin-1 as substrate and converts it into a dienone amide was isolated and characterized from *A. cavernicola* ^27^.

Despite being heavily chemically defended, *A. aerophoba* is grazed by the sea slug *T. perversa*. While feeding, this sea slug selectively sequesters and accumulates sponge-derived brominated alkaloids, in particular aerophobin-2 ^28^. It also accumulates *A. aerophoba’s* saffron-coloured pigment uranidine, which makes *T. perversa* camouflage with the sponge ^29^. We performed controlled experiments to characterize the response of *A. aerophoba* to grazing by *T. perversa*. To further elucidate if the sponge response is specific to grazing by this specialist, we included a treatment in which sponges were mechanically-damaged with a scalpel. At the cellular level, we focused on the fate of spherulous cells, in which brominated compounds are stored ^30^, via light and transmission electron microscopy. At the chemical level, we visualized the distribution of two of the main brominated compounds in *A. aerophoba*, aerophobin-2 (precursor) and aeroplysinin-1 (bioconverted product) ^24,25^ by Matrix-Assisted Laser Desorption-Ionization imaging Mass Spectrometry (MALDI-imaging MS), in order to avoid bias derived from chemical extraction procedures. Under the hypothesis of bioconversion, we expected to detect higher concentrations of aeroplysinin-1 in wounded than in control samples, and the opposite pattern for aerophobin-2. We further tested whether the chemical and cellular responses translated into increased deterrence capability against *T. perversa*.

## METHODS AND MATERIALS

### Animal Collection

The sponge *Aplysina aerophoba* and the sea slug *Tylodina perversa* were collected by Scuba diving in June 2016 and May 2017 at the Mediterranean coast of Spain (42.29408 N, 3.28944 E and 42.1145863 N, 3.168486 E, respectively), at a depth between 2 to 10 m. Animals were immediately transported to the Experimental Aquaria Zone at the Institute of Marine Science (ICM-CSIC) in Barcelona (Spain). There, each sponge individual was divided into 2-3 specimens with its own osculum (**Fig. 1**). Each specimen was placed into individual 6L aquaria and maintained in a flow-through system with direct intake of seawater.

**Figure 1.**
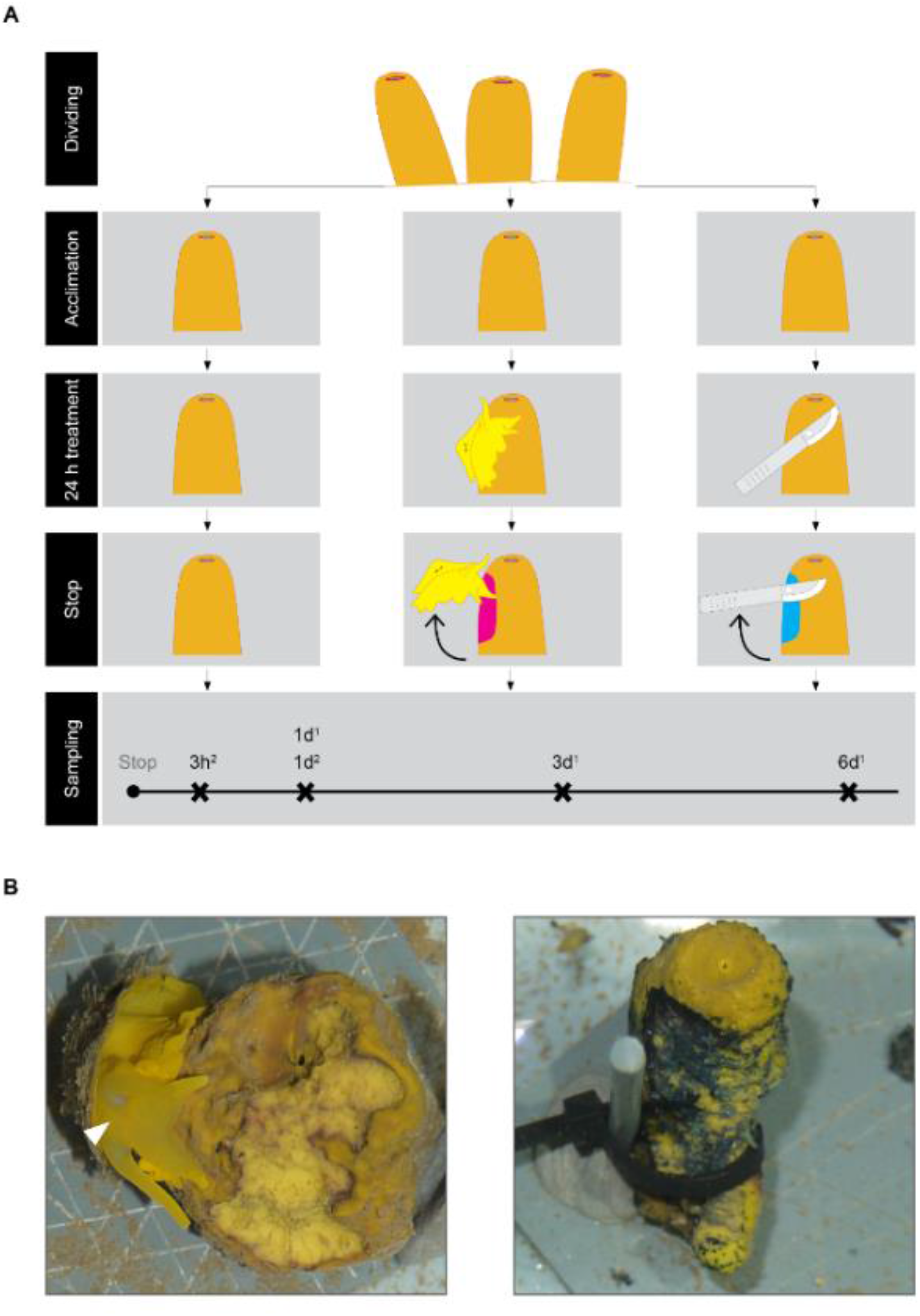
Experimental design. (**A**) Each sponge individual was divided into three specimens that were randomly assigned to either control (left panels), grazing (middle panels), or mechanical damage (right panels) treatment. Treatment was applied for 24 hours. We performed consecutive experiments to collect samples at different time points after stop of treatment: 3 h, 1 day, 3 days, and 6 days. Superscript number 1 and 2 denotes experiments performed in 2016 and 2017, respectively. (**B**) Wound by *T. perversa* (arrowhead, left panel) and by mechanical damage (right panel).

### Experimental Set-up

After 1 week acclimation, the specimens from each sponge individual were randomly assigned to one of the following treatments (i) control: no treatment, (ii) grazing: one sea slug, that had been starved for 24 h, was placed in direct contact to the sponge specimen and allowed to feed *ad libitum* for 24 h, and (iii) mechanical damage: the sponge specimen was clipped at the surface with a scalpel for 3 min every half hour for the first 3 hours and the last 3 hours of the 24h-treatment period (**Fig. 1**). All treatments were stopped after 24 h. Each experiment had the same experimental design but differed in the sampling point after stop of treatments (**Fig. 1**): three experiments were performed in 2016, in which samples were collected 1 day, 3 days, and 6 days after stop of the treatment; two experiments were performed in 2017 and samples were collected 3 h and 1 day after stop of the treatment (n = 3-4 for all experiments, **Supplementary Table 1 online**). No mechanical damage group was performed in the 3-day experiment.

### Sample Preparation for Light Microscopy

Samples were immediately fixed in 2.5 % glutaraldehyde in 0.1 M cacodylate buffer/11 % sucrose and stored at 4 °C. Samples were then processed as in Wu, 2019 ^31^. Semi-thin (0.35 μm) sections were prepared with an ultramicrotome (Reichert Ultracut S, Leica, Austria), deposited on Superfrost Plus glass slides (Menzel) with Biomount 2 mounting medium (BBI Solutions), stained with Richardson solution, and imaged (100x magnification) with a ZEISS Axio Observer microscope (version 1.1, Zeiss, Germany).

### Automatic Counting of Spherulous Cells

Microscopic images were analysed in ImageJ (version 1.51j8, Java 1.8.0_112) ^32^. The first 100 μm from the surface to the inside was defined as the Region Of Interest 1 (ROI 1, **Supplementary Fig. 1 online**) and its area was measured excluding aquiferous canals. Image type, Substract background, Threshold, and Watersheld parameters were adjusted to select densely-stained spherulous cells. Next, those cells were automatically counted by using “Analyze Particles” ImageJ tool with “Size” = 34 - 314 μm^2^ (considering round cells with 10 - 20 μm of diameter and a sectioning at 1/8 from cell edge or through cell center) and “Circularity” = 0.20 - 1.00 (considering cell shapes from elongated/possibly motile stage to circular/possibly non-motile stage). Spherulous cells on edges of each ROI were not counted. The number of densely-stained spherulous cells pro 50 000 μm^2^ area was calculated for the ROI 1 of each image (**Supplementary Fig. 1 online**) compared amongst treatments by applying Generalized Linear Mixed-effects Model via Penalized Quasi-Likelihood (glmmPQL) in R (v3.6.0) ^33^ as implemented in RStudio (v1.2.1335) ^34^, with treatment as the fixed effect and sponge individual as the random effect. The distribution pattern of the densely-stained spherulous cells from the surface to the inside of sponges was investigated by defining 5 consecutive 100 μm-deep ROIs adjacent to ROI 1 (**Supplementary Fig. 1 online**) and counting cells following the methodology described above. The automatic counting was validated by manual counting on a subset of samples (**Supplementary Fig. 1 online)**.

### Transmission Electron Microscopy (TEM)

One sponge individual from 1d, with its corresponding three differently-treated specimens, was analysed by TEM in order to characterize the spherulous cells observed by light microscopy. Embedded samples were cut into ultra-thin (70 nm) sections using an ultramicrotome (Reichert Ultracut S, Leica, Austria). Sections were mounted on pioloform coated grids and contrasted with uranyl acetate replacement stain (Science Services, Germany) for 20 min and subsequently with Reynold’s lead citrate for 3 min. Ultra-thin sections were imaged with a Tecnai G2 Spirit BioTwin transmission electron microscope (80 kV, FEI, USA) at the Central Microscopy of University of Kiel (Germany).

### Sample Preparation for MALDI-imaging MS

The spatial distribution of secondary metabolites within sponge cross-sections was assessed by MALDI-imaging MS. Samples from 1d-2017 experiment (*n*=4 individuals x 3 treatments) were wrapped in aluminium foil (with a clear annotation of the wound location in grazing and mechanical damage treatments), snap-frozen in liquid nitrogen, and immediately stored at −20 °C. Samples were prepared as described by Yarnold *et al*. ^35^, with some modifications (see details in **Supplementary Information online**). In short, each sample was frozen-sectioned at 14 μm and mounted onto Indium-Tin-Oxide (ITO, Bruker Daltonics, Bremen, Germany). Each ITO glass slide consisted of three sections corresponding to three sponge specimens (control, grazed, and mechanically-damaged) from the same sponge individual. Unlike Yarnold et al. ^35^, the sample sections were directly mounted onto the slides without previous washing with MilliQ-water, as this step caused a morphological alteration of our sections resulting in the delocalization of the compounds of interest. After light microscopy imaging (100x magnification), each ITO glass slide was coated with universal MALDI matrix chosen for optimal visualisation of both aerophobin-2 and aeroplysinin-1, which differ in polarity. Further information on the optimization of the sample preparation process can be found in Wu, 2019 ^31^.

### MALDI-imaging MS Analysis

In MALDI-imaging MS, a laser is passed over the sample and, due to the matrix, compound ions are released and passed to the mass analyser. At each raster point, mass spectrometry data is obtained and then integrated into an image in a specialized software. Here, raster widths of 250, 275, and 300 μm were selected according to the section area and in order to preserve sensitivity and consistency during measurement (**Supplementary Table 2 online**). Samples were analysed in an UltrafleXtreme MALDI TOF/TOF (Bruker Daltonics), operated in positive reflector (detailed analysis is provided in **Supplementary Information online**).

Brominated compounds with two bromine atoms (Br) show a specific three-peak-pattern because the two stable isotopes (^79^Br and ^81^Br) occur in similar abundance in nature (ratio 50.5 : 49.5). Thus, aerophobin-2 and aeroplysinin-1 were identified by their molecular mass and isotopic pattern. Commercially available aerophobin-2 and aeroplysinin-1 (Santa Cruz Biotechnology, Germany) served as standard references to investigate ionization yields at the same concentration (1 mM). Since aerophobin-2 and aeroplysinin-1 showed different ionization yields, relative intensity of each compound between treatments was qualitatively compared. First, MALDI-imaging MS datasets were Root Mean Square normalised. Then, the intensity for each compound was shown as relative intensity to the highest value among the three sections within each ITO glass slide (i.e., among the specimens of the same individual), and depicted in a colour scale.

We investigated the co-localization of aerophobin-2 and aeroplysinin-1. Ion images were exported as greyscale images in SCiLS Lab 2016b so that in each pixel, the absolute intensity of each compound was computed as relative values of a gradient grayscale (0 to 255). The resulting images were imported in Fiji ^36^ and analysed as follows: Co-localization images show the pixels where both compounds had a value >0 in the greyscale (abbreviated AND). Occurrence images show the pixels where at least one of the two compounds had a value >0 in the greyscale (abbreviated OR).

MALDI-imaging MS allows the untargeted monitorisation of *A. aerophoba* metabolites, with thousands of complex spectra per sample. Compounds containing Br were manually annotated according to their isotopic profile. Moreover, the whole datasets were explored by a multivariate statistical tool, spatial segmentation: Spectra were clustered by their similarity, generating an unsupervised spatial segmentation map of each tissue section where regions of similar chemical compositions (i.e. clusters) were revealed and depicted with a distinct colour ^37^. Segmentation maps were calculated in SCiLS Lab 2015b by bisecting k-means using correlation distance, weak denoising and Root Mean Square normalisation (peak picking workflow: 200 peaks (every 2 spectra).

### Deterrence experiments

We tested if *T. perversa* preferred control over mechanically-damaged sponges. In the 2017 animal collection effort, we took one chimney of five individuals of *A. aerophoba* and five individuals of *T. perversa*. We ran the experiment one day after collection, following suggestions in ^38^. One hour before the experiment, each sponge chimney was divided into apical, middle, and bottom portions. Each portion was separated through the middle of the osculum (ca. 3 to 6 cm^3^) into two pieces which were randomly assigned to either control or mechanical damage treatment. Mechanical damage was applied during 5 min and the wound was 1 cm^2^ and 1-2 mm deep. To test deterrence, we covered *T. perversa* with an opaque vessel for 15 min. We placed two pieces of the same sponge individual and region, one mechanically-damaged and one control, at 5 cm of the sea slug and we removed the vessel. We considered that the sea slug made a choice when it touched the sponge with the oral tentacles or with the front head within the first 15 min after removal of the vessel (**Supplementary video 1 online**). *T. perversa* preference was tested in a Binomial test (p=q=0.5) in R (v3.6.0) ^33^ as implemented in RStudio (v1.2.1335) ^34^.

## RESULTS

Once the experiment started, each sea slug usually took 15 – 30 min to start feeding on the sponge (video at doi.pangaea.de/10.1594/PANGAEA.907958). They tightly attached and covered the sponge with the mantle during grazing. The oscula of control group sponges were usually wide open, whereas they were less open or even closed in grazing or mechanical damage group sponges. Sea slugs usually remained at the same spot on the sponge during the 24h feeding period, turning over their bodies to feed neighbour tissue. The wounds generated by grazing showed a bright yellow colour, similar to the ones observed in the field, whereas the wounds in mechanical damage were dark blue (**Fig. 1**). When grazing-caused wounding showed some darkening, it was comparatively minor and occurred at the frontier between the wound and the intact tissue.

### Accumulation of Spherulous Cells at the Wound

Images (100x magnification) of specimens collected 1d after wounding showed a striking accumulation of densely-stained cells at the injured surface (first 100 μm from the wound to the inside) in both grazed and mechanically-damaged specimens but not in the controls (**Fig. 2A**). Further inspection by TEM confirmed that those cells were spherulous cells with electron-dense spherules (termed spherulous cells for the following text), (**Fig. 2B**). In contrast, the surface of the control group contained mostly spherulous cells with electron-lucent spherules (**Fig. 2C**). In addition, shedding of spherulous cells and the presence of cell debris at the wound site and the aquiferous canals occurred more frequently in both grazed and mechanically-damaged specimens than in the control (**Supplementary Fig. 2 online**).

**Figure 2.**
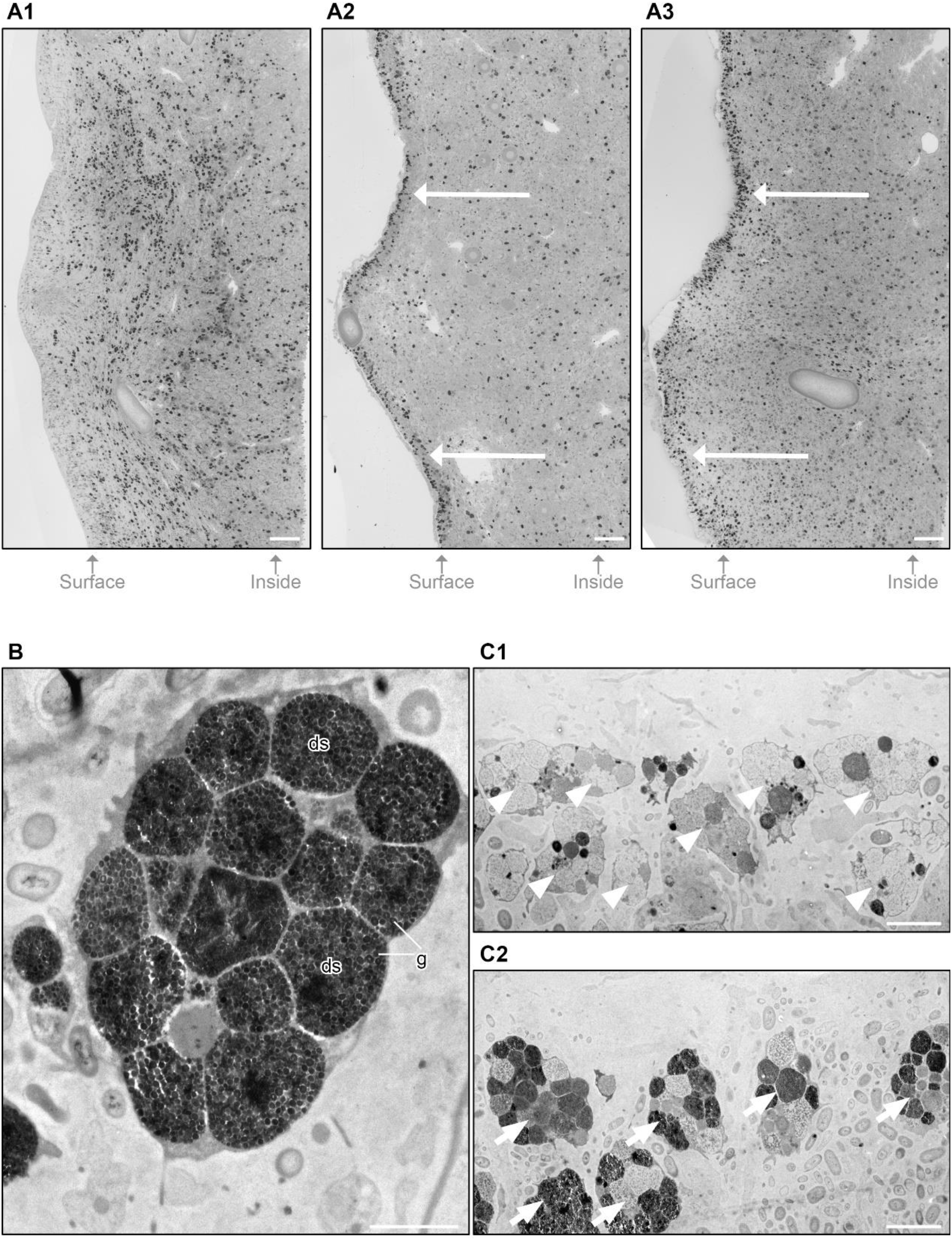
Accumulation of spherulous cells with electron-dense spherules at the surface 1d after stop of treatments. **(A)** Microscopic section (100x magnification) of 1d-samples showing that densely-stained spherulous cells accumulated at the wound (arrow) in grazed (A2) and mechanical damage (A3) groups compared to control (A1). Scale bar= 100 μm. (**B**) TEM-image at the wound confirming that densely-stained spherulous cells correspond with spherulous cells containing electron-dense spherules (ds), with numerous electron-dense granules (g). Scale bar= 2 μm. (**C**) The surface of control group contained spherulous cells with electron-lucent spherules (C1, arrowhead). The surface of wounding group contained spherulous cells with electron-dense spherules (C2, arrow). Scale bar= 5 μm.

We further quantified the accumulation of spherulous cells at different time points. Upon grazing, spherulous cells gathered at the surface 3h after treatment (glmmPQL, *p* = 0.023). After 1d, the density of spherulous cells at the surface reached the highest value in both the grazed and mechanically-damaged groups (**Fig. 3A**) and represented a significant accumulation compared to control samples, except in 1d-2016 grazing group (glmmPQL; 1d-2016: grazing, *p* = 0.056 and mechanical damage, *p* = 0.022; 1d-2017: grazing, *p* = 0.035 and mechanical damage, *p* = 0.013). In contrast, 3-d and 6-d samples showed a similar density of spherulous cells at the surface amongst all treatments (glmmPQL, *p* > 0.1).

**Figure 3.**
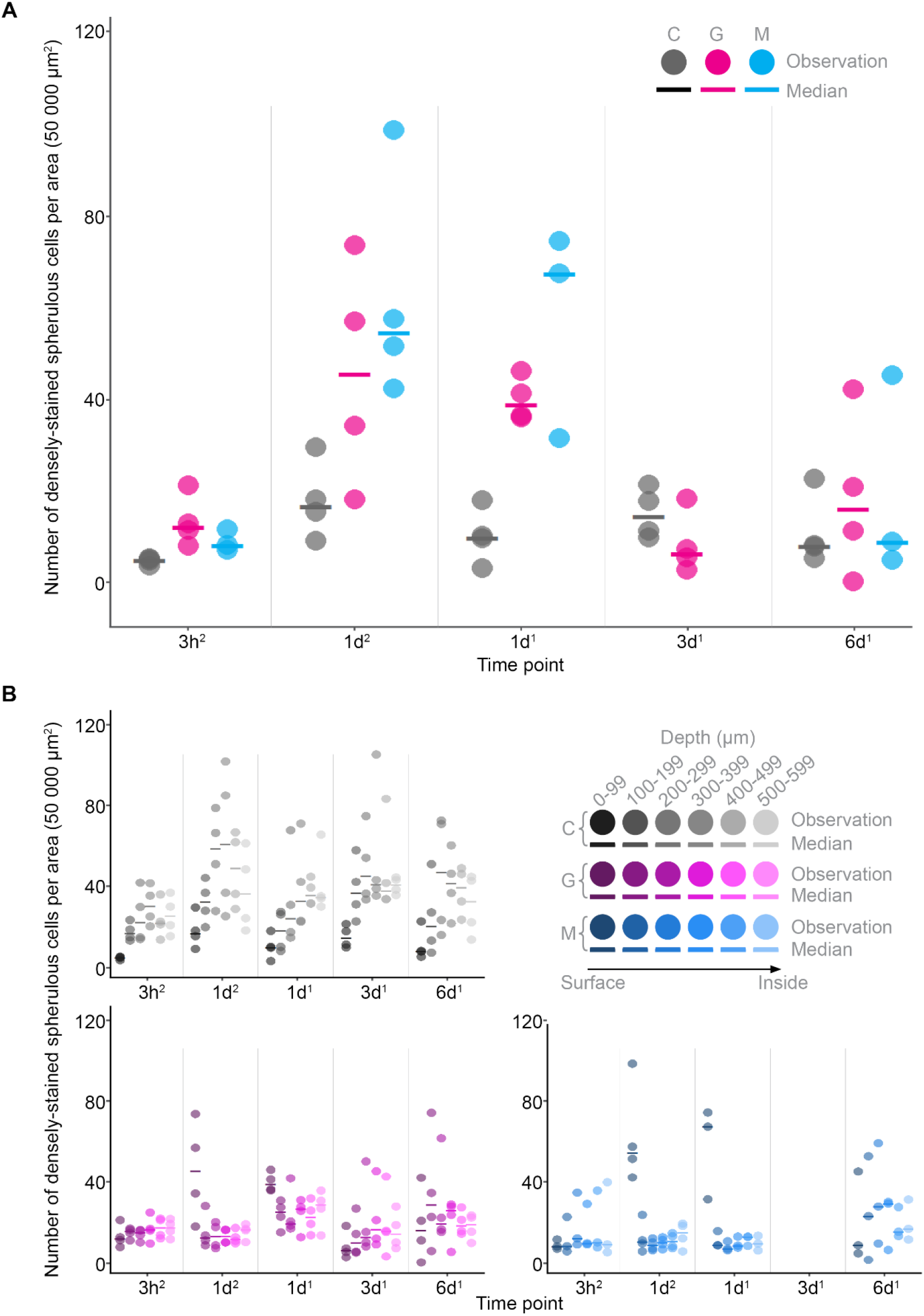
Time-dependent re-distribution of spherulous cells upon wounding. Number of densely-stained spherulous cells per area (50 000 μm^2^) right at the surface (first 100 μm) (**panel A**) and within the first 600 μm (**panel B**). Note that there was no mechanical damage group at 3d. Superscript number 1 and 2 denotes experiments performed in 2016 and 2017, respectively. C=control; G=grazing; M=mechanical damage

We also investigated the distribution of spherulous cells from the surface to 600 μm inside the tissue. In the control group, the density of spherulous cells followed a depth-dependent distribution pattern with a lower value at the surface (first 100 μm) compared to the inner tissue, with the highest value at a depth of ca. 300 - 400 μm (**Fig. 3B**). After grazing and mechanical damage, this distribution pattern was disrupted in both 3h and 1d samples (**Fig. 3B**). At 3h after wounding, sponges displayed a similar density of spherulous cells from the wounded surface to the inside of the sponge specimens. At 1d, the cell density peaked at the wounded surface, whereas the density in the inner tissue was similar to that observed at 3h. After 3d, the distribution pattern of spherulous cells resembled that observed in control group, consistent with the initiation of regeneration (**Supplementary Fig. 3 online**). In fact, after 3 days, the ectosome showed a well-defined surface border with less densely-stained spherulous cells (**Supplementary Fig. 3 online**).

### High Inter-individual Variability of the Distribution of Brominated Alkaloids but Distinct Allocation upon Wounding

We visualised the abundance of two of the main brominated secondary metabolites in *A. aerophoba*, aerophobin-2 and aeroplysinin-1, within the 1-d after treatment sponges by MALDI-imaging MS. Both aerophobin-2 and aeroplysinin-1 occurred in all samples (experimental monoisotopic *m/z* 504.0 corresponding to C_16_H_20_Br_2_N_5_O_4_ for aerophobin-2, and monoisotopic *m/z* 337.7 corresponding to C_9_H_10_Br_2_NO_3_ for aeroplysinin-1). These isotopic patterns agreed with their molecular formula and matched the isotopic patterns of the standards (**Fig. 4A**). MALDI-imaging MS revealed other Br-containing compounds in all samples, the ones with higher abundance are shown in **Supplementary Figure 4 online**. The putative molecular formula associated to these isotopic patterns pointed to brominated alkaloids. These detected metabolites might be those already reported in *Aplysina* species (e.g. monoisotopic experimental *m/z* 420.954, putative molecular formula [M+H]^+^ C16H27Br2N2O fits with the molecular formula of Aplysamine-1) or related to other known *Aplysina* brominated alkaloids (e.g. most intense isotope experimental *m/z* 1138.352, putative molecular formula [M+H]^+^ C_33_H_31_Br_6_N_4_O_11_ is most likely related to fistularins such as fistularin-3 C_31_H_30_Br_6_N_4_O_11_) (**Supplementary Fig. 4 online**).

**Figure 4.**
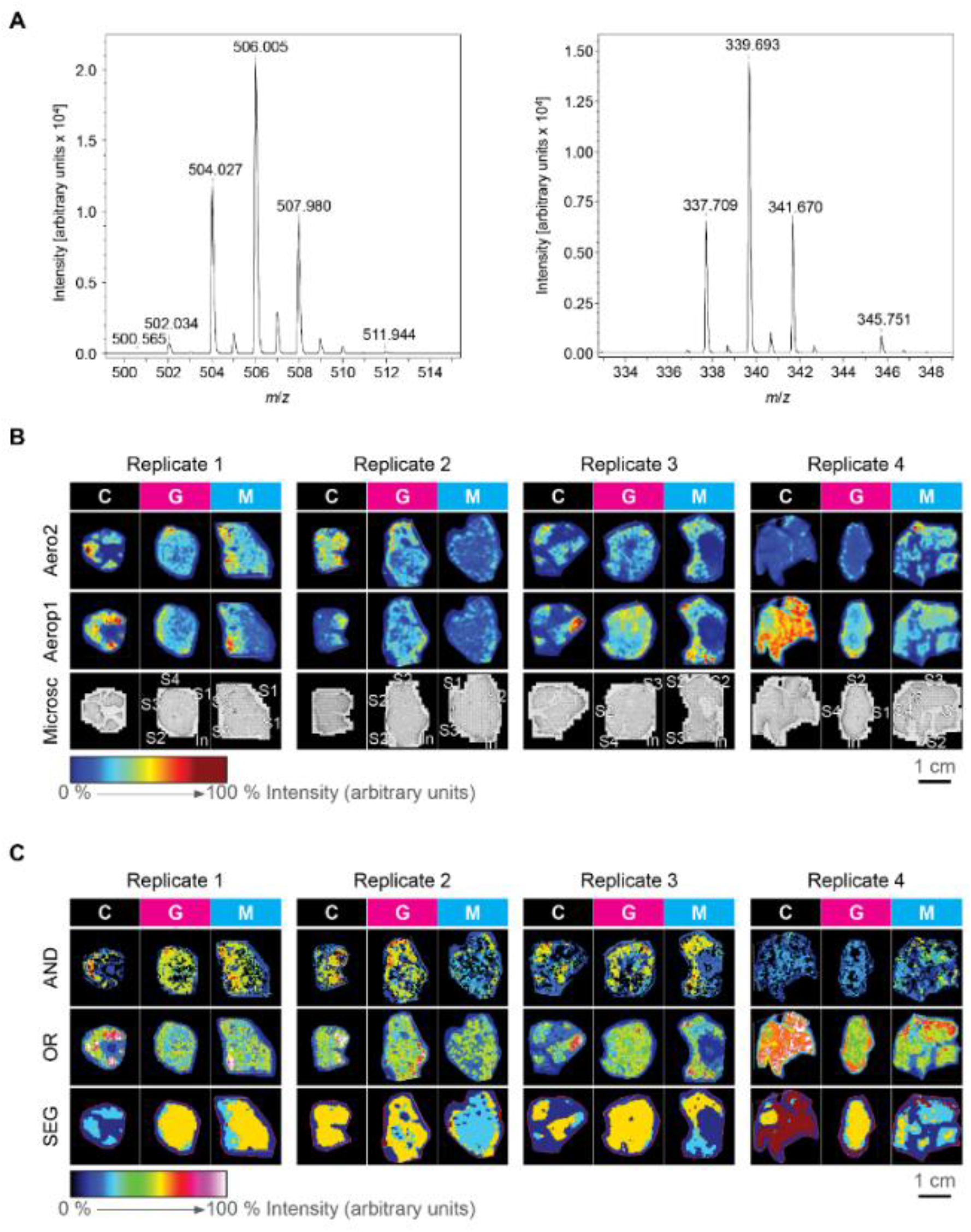
MALDI-imaging MS of sponges 1d after stop of treatments. The experimental isotopic patterns of aerophobin-2 (left panel) and aeroplysinin-1 (right panel) by MALDI-imaging MS. (**B**) The relative abundance of these compounds (Aero2=aerophobin-2, Aerop1=aeroplysinin-1) in the three different treatments (C=control, G=grazing, and M=mechanical damage) for each biological replicate (Replicate; i.e., specimens of a same sponge individual). The corresponding microscopic images (Microsc) are also shown. (**C**) Images of the co-localization of aerophobin-2 and aeroplysinin-1 (AND), the distribution of at least one of these two compounds (OR), and the segmentation map (SEG). The relative intensity from 0 to 100 % is depicted in a colour scale with warmer colours representing relatively higher intensity and colder colours lower intensity of each compound. Scale bar = 1 cm; In= intact surface; S1= the first evident wound; S2= the second evident wound, and so on; White-dotted line= broken or cut edges.

MALDI-imaging MS revealed a striking biological variability of the distribution of both aerophobin-2 and aeroplysinin-1 among the four control samples (**Fig. 4B**). Interestingly, both grazing and mechanical damage resulted in a re-distribution of these two compounds in a way that was different from the control and more consistent among the biological replicates, with preferential accumulation of both aerophobin-2 and aeroplysinin-1 at the surface (**Fig. 4B**). Unsupervised segmentation maps based on the 200 most abundant compounds clustered together (colour coded) regions of the sample with specific molecular composition ^37^ and confirmed the distinct patterning in wounding groups (**Fig. 4C**). Segmentation maps from control specimens were, in general, chemically homogeneous (and thus control sections are mostly represented by one colour), while treated specimens showed chemical differences between the surface and the inner part of the sponge (and thus are represented by two colours) (**Fig. 4C**). The distribution of unknown brominated compounds followed a similar pattern as aerophobin-2 and aeroplysinin-1 (**Supplementary Fig. 5 online**).

The results showed two consistent patterns in the distribution of aerophobin-2 and aeroplysinin-1 among all control, grazed and mechanically-damaged samples (**Fig. 4C**): (i) the images showing the co-localization of the two compounds (**Fig. 4C, AND**) correlated with the spatial distribution of aerophobin-2; and (ii) areas with the highest intensity in the occurrence maps showing at least one of the two compounds (**Fig. 4C, OR**) usually corresponded to no signals in the co-localization images (**Fig. 4C, AND**) indicating that only one of these compounds accounts for the signal in those areas. These spatial associations suggest that: (i) aeroplysinin-1 correlated with aerophobin-2 (i.e., when aerophobin-2 is detected, aeroplysinin-1 is usually detected as well); and (ii) a higher intensity of one compound was coincident with decreased intensity of another compound, which suggests interconversion. These two phenomena took place independently from the treatments.

### Correlation between Brominated Alkaloids and Spherulous Cell Accumulation

The different resolution of the microscopy images used for automatic counting and the MALDI-imaging MS images prevents us from unequivocally assigning the accumulation of brominated alkaloids to the accumulation of spherulous cells at the wound. In MALDI-imaging MS technology, we ran the samples at a resolution of 250-300 μm to ensure the comparability of the within-tissue distribution of target compounds among the different treatments, while keeping the experiments short enough to maintain constant intensity **(Fig. 4**). However, we did analyse one grazed sponge collected from the field at higher spatial resolution (20 – 100 μm) and we observed that a “track” of spherulous cells co-localized with peaks of both aerophobin-2 and aeroplysinin-1 (**Fig. 5**).

**Figure 5.**
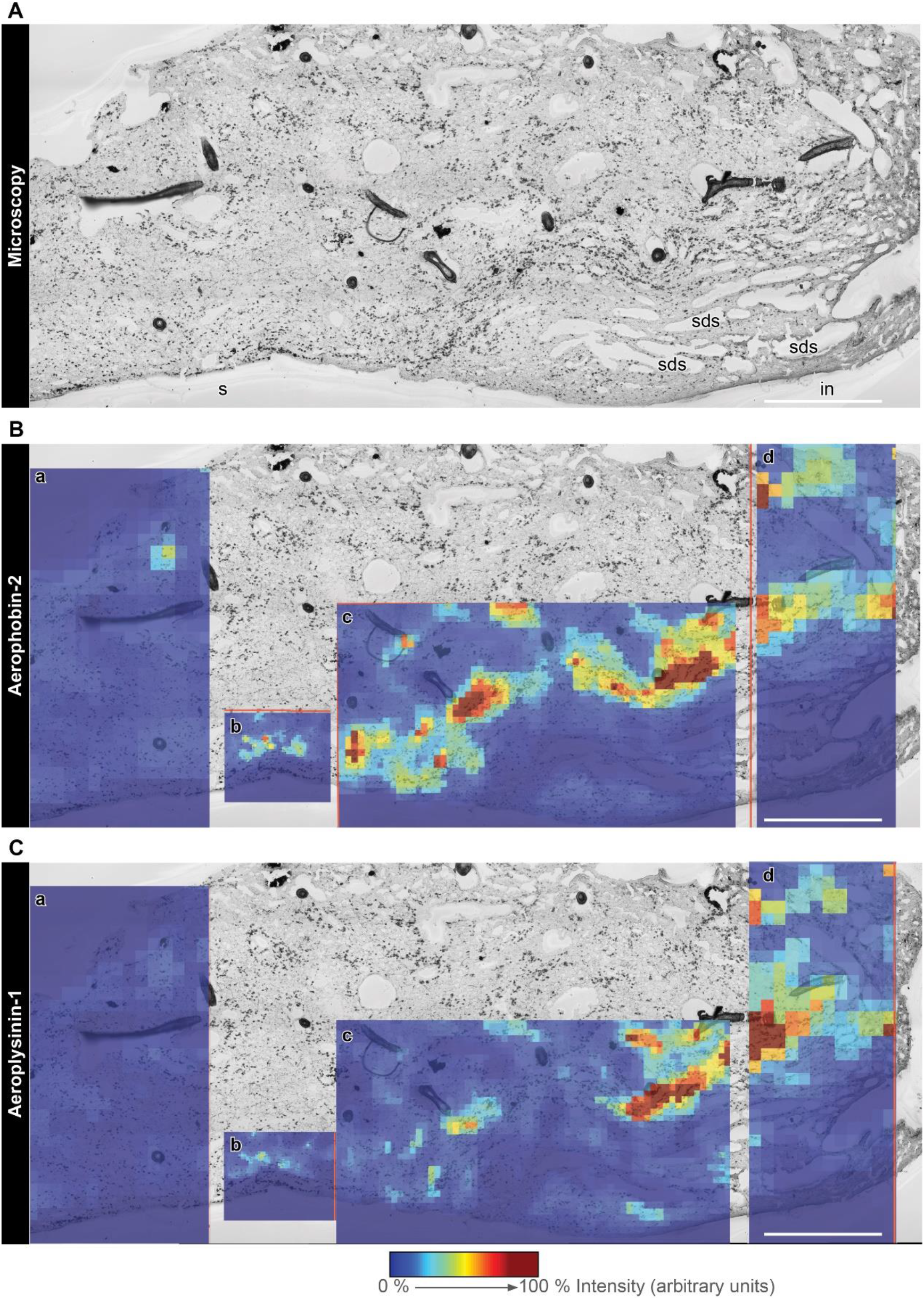
Correlation of cellular pattern and spatial distribution of brominated compounds. (**A**) Light microscopic image of a grazed sample collected from the field. A track of spherulous cells was parallel to the surface beneath the subdermal spaces (sds) below the intact surface (in) and at the injured surface (s). Superimposition of the microscopic image with 2D-MALDI-images of aerophobin-2 (**B**) and aeroplysinin-1 (**C**), respectively. The sample was measured by MALDI-imaging MS with a raster size at 100 μm (a and d), 50 μm (c), and 20 μm (b). The relative intensity of each compound is depicted in a colour scale. Scale bar = 1 mm.

#### <u>Tylodina perversa</u> prefers non-damaged sponges

In the deterrence experiments, if a choice was made (**Supplementary Video 1 online**), *T. perversa* always preferred control sponge pieces over mechanically-damaged ones (**Table 1**; Binomial test with p=q=0.5, P= 0.002). One *T. perversa* individual remained still in all experiments and made no choice. This individual was bigger than the others and showed less mobility. The experiment included different portions of chimney to reduce potential intra-individual variability and we observed fewer choices in the experiments including the middle part of the sponge (**Table 1**).

**Table 1.** Feeding choice experiments in *Tylodina perversa*. The total number of replicates is expressed by “N”. “No choice” denotes those experiments where *T. perversa* did not show a preference within 15 min after the sponges were presented. “Choice” shows the preference and number of times that option was selected.

| Feeding choice | N | No choice | Choice |
| --- | --- | --- | --- |
| Control : Mechanical Damage | 15 | 6 | 9:0 |
| Chimney portion - Apical :Middle : Bottom |  |  | 4:1:4 |
| *T. perversa* individual- 1:2:3:4:5 |  |  | 2:0:3:2:2 |
| *A. aerophoba* individual- 1:2:3:4:5 |  |  | 2:1:2:2:2 |

## DISCUSSION

Our results show that the response of *A. aerophoba* to grazing by the specialist *T. perversa* resembles the response to a general wounding (here the mechanical damage treatment). MALDI-imaging MS showed that the brominated alkaloids aerophobin-2 and aeroplysinin-1 are constitutively present in all sponge samples and their distribution within the tissue varies among individuals. Yet, these secondary metabolites were consistently re-allocated to the surface upon wounding. Spherulous cells, carrying brominated alkaloids, gathered at the wound within the first day after wounding. After 3d, coincident to visible signs of tissue restoration, all treatments showed similar spherulous cell distribution.

We propose that the active migration of spherulous cells plays a role in defence and regeneration upon wounding. Previous studies ^30^ and the MALDI-imaging MS results at high spatial resolution (**Fig. 5**) support their function as carriers of the brominated alkaloids. The sponge *Crambe crambe* exudes toxic compounds via the release of spherulous cells to the exhalant water of the sponge upon mechanical stress in a process called spherulisation ^39^. We also observed the shedding of spherulous cells upon damage and their release into aquiferous canals; thus, spherulisation may be also occurring in *A. aerophoba*. Recently, Ereskovsky et al. ^40^ described the accumulation of spherulous cells at the wound, 1 day after applying a small excision, as part of the regenerative blastema in *Aplysina cavernicola. A. aerophoba* showed a similar cellular response, peaking also after 1 day, in both mechanical damage and grazing treatments, even if in our study wounding damage was stronger than in Ereskovsky et al.^40^. Therefore, the mobilisation of spherulous cells may be a common process involved in sponge defence, but also provide structure or energy for rapid regeneration.

Our results suggest that the bioconversion between the precursor aerophobin-2 and aeroplysinin-1 is constitutive and occurred in all samples, including control as well as wounding treatments. We did observe that both compounds were less abundant in treated samples than in controls. One explanation could be their transformation to the related dienone, which we could not detect in our MALDI-imaging MS protocol, likely because dienone signal overlaps with background matrix signals. Another explanation could be their release to the environment via spherulisation. It is remarkable that the different distribution between control and treated samples not only affects aerophobin-2 and aeroplysin-1, but also other compounds (as suggested by segmentation maps in **Fig. 5C**) and other brominated compounds (**Supplementary Fig. 4 online**).

MALDI-imaging MS revealed the dynamic nature of the sponges’ chemical response to predation and wounding with unprecedent spatial resolution. We observed high variability in the distribution of chemical compounds in control specimens, in concordance with the natural variability in absolute concentrations observed in previous chemical studies of *A. aerophoba* ^18,41,42^, and other sponge species (reviewed in ^12^). Several studies explored the differential allocation of secondary metabolites in sponges, with concentrations usually higher in the inner than the outer parts of the sponges ^12^. This pattern may reflect the antibacterial activity function of secondary metabolites ^16,43^. The numerous aquiferous canals in the inner sponge tissue effectively expand the surface area of the sponge exposed to the external environment and Optimal Defence Theory predicts that defences should be allocated in those regions that are most valuable and/or at higher risk. However, our results did not show a clear differential allocation but rather strong individual variability in control samples. This variability contrasted to the consistent gathering of compounds to the sponge surface upon wounding; a pattern which seems to arise from a reallocation of existing defences rather than *de novo* synthesis. Such reallocation would have the advantage of using available resources to enhance the protection of the area that has now been signalled as most vulnerable to damage, fitting the postulates of Optimal Defence Theory. Moreover, this mechanism would provide a flexible use of chemical defences to adapt to different challenges and may explain the recurrent intraspecific variability in sponges.

Despite the similarity in the response upon grazing and upon mechanical damage, we did detect one prominent difference between the two treatments. Upon mechanical damage, sponge wounds turned black due to the oxidation of uranidine, which likely happens during cell damage or exposure to the surrounding environment ^29^. Our experiments suggest that this reaction deters *Tylodina* and, in fact, wounds in the grazing treatment remained mainly yellow. *Tylodina* grazes with its radula in a way that the wound remains covered by its mantle and/or by the mucus liberated by the slug during feeding. This may serve to reduce cellular damage and exposure of the wound to the environment; thus, preventing the formation of more deterrent sponge compounds.

Grazing by the specialist sea slug *T. perversa* triggers a wound-like response in the sponge *A. aerophoba*. This response consisted of a local accumulation of spherulous cells, which are likely directed to enhanced regeneration ^40^ as well as defend the exposed wound against invading microbes and predators ^25^. After one day, coinciding with the peak of spherulous cell accumulation, brominated secondary metabolites are re-allocated to the surface of the sponge. We propose that wounding cues signal the surface as the region at most risk and thus, induce reallocation. We observed the darkening of the sponge wound immediately upon mechanical damage, which is likely due to the release of oxidized compounds ^29^. Interestingly, *Tylodina* prevents this from happening, probably because it covers the wound with its mantle while feeding. Thus, it seems that, contrary to reports in other organisms such as algae or dinoflagellates ^44–46^, the sponge has not evolved a specific response to the specialist grazer and rather relies on the rapid regeneration and reallocation of existing resources as key strategies for coping with predation and wounding.

## DECLARATIONS

### Availability of data and material

Images derived from light microscopy and MALDI-imaging MS and videos recorded during the experiments are available at PANGAEA, https://doi.org/10.1594/PANGAEA.907958

### Author contribution statement

Y.W., L.P. and U.H. conceived the idea. Y.W., B.P., L.P. and M.R. planned and conducted the experiments. B.P. analysed the deterrence experiment data. Y.W. performed and analysed the microscopy and prepared the samples for MALDI-imaging MS. M.G.-A. ran and analysed MALDI-imaging MS. Y.W. and L.P. wrote the manuscript. All authors made substantial contribution to the writing of the manuscript and approved it for publication.

## Acknowledgements

We thank Rafel Coma, Manel Bolívar and Marc Catllà (CSIC) and Laura Rix (University of Queensland) for assistance during sponge experimentation. We thank the “Parc Natural del Montgrí, les Illes Medes I el Baix Ter” and “Parc Natural del Cap de Creus” for sampling permissions, Martin Wahl and Mark Lenz (GEOMAR) for statistical comments, and the Central Microscopy Facility (University of Kiel) for microscopy support.

