## Supplementary tables and figures for "Grazing results in mobilization of spherulous cells and re-allocation of secondary metabolites to the surface in the sponge *Aplysina aerophoba*"

### Supplementary Information

**Protocol for MALDI-imaging MS sample preparation and analysis.** Samples were cryosectioned as described by Yarnold *et al.* (2012), with some modifications. In short, each sample was thawed at RT and placed in a cryomold (22 mm in diameter, 5 mm in height, Tissue-Tek@ Cryomold@, Plano, Germany) with a drop of embedding medium, Optimal Cutting Temperature (OCT, Tissue-Tek@, Plano, Germany). If samples were bigger than the cryomold, exceeded sponge tissue was cut to fit the cryomold, preserving the surface. After filling with OCT, each sample was frozen-sectioned at 14 µm in a cryostat (CM3050 S, Leica, Germany) at -20 °C and then thaw-mounted onto Indium-Tin-Oxide (ITO, Bruker Daltonics, Bremen, Germany) for subsequent MALDI-imaging MS. Each ITO glass slide was spray-coated with 2 ml of a saturated solution (20 mg/mL) of universal MALDI matrix (1:1 mixture of 2,5-dihydroxybenzoic acid and α-cyano-4-hydroxy-cinnamic acid; Bruker Daltonics, Germany) in acetonitrile/methanol (70:30, v/v), using the automatic system ImagePrep device 2.0 (Bruker Daltonics) in 60 consecutive cycles (the sample was rotated 180° after 30 cycles) of 41 seconds (1 s spraying, 10 s incubation time, and 30 s of active drying). The matrix solvent selection was optimized to visualize the spatial distribution of both aerophobin-2 and aeroplysinin-1, which differ in polarity.

Samples were analysed in an UltrafleXtreme MALDI TOF/TOF (Bruker Daltonics), operated in positive reflector mode using flexControl 3.0. The analysis was performed in the 100-1500 Da range and with 30 % laser intensity (laser type 4), accumulating 1000 shots by tanking 50 random shots at every raster position. External calibration of the acquisition method was performed using Peptide Calibration Standard II (Bruker Daltonics) containing Bradykinin1-7, Angiotensin II, Angiotensin I, Substance P, Bombesin, ACTH clip1-17, ACTH clip18-39, and Somatostatin 28. Spectra were processed with baseline subtraction in flexAnalysis 3.3 and corrected internally using the peaks of HCCA ($[M+H]^+$ *m/z* 190.0499 and $[2M+H]^+$ *m/z* 379.0925). MALDI-imaging MS data was visualised in SCiLS Lab 2015b (SCiLS, Bremen, Germany).

**Supplementary Table 1. Number of biological replicates for microscopy and MALDI-imaging MS. C**: control; **G**: grazing; **M**: mechanical damage.

| | Microscopy | | | MALDI-imaging MS | | |
|---|---|---|---|---|---|---|
| **Experiment** | **C** | **G** | **M** | **C** | **G** | **M** |
| **3h-2017** | 4 | 4 | 3 | | | |
| **1d-2017** | 4 | 4 | 4 | 4 | 4 | 4 |
| **1d-2016** | 4 | 4 | 3 | | | |
| **3d-2016** | 4 | 4 | | | | |
| **6d-2016** | 4 | 4 | 3 | | | |
| **Total** | | 53 | | | 12 | |

**Supplementary Table 2. Used raster size, measured area, and number of mass spectra acquired by MALDI-imaging MS from 1-d sponge specimens.** Individual: sponge individual; C: control; G: grazing; M: mechanical damage.

| Individual | Treatment | Raster [$\mu m^2$] | Area [mm²] | Number of spectra |
|---|---|---|---|---|

| | | | | |
|---|---|---|---|---|
| 1 | C | 250 x 250 | 131.06 | 2097 |
| 1 | G | 250 x 250 | 167.06 | 2673 |
| 1 | M | 250 x 250 | 233.44 | 3735 |
| 2 | C | 275 x 275 | 131.29 | 1736 |
| 2 | G | 275 x 275 | 216.89 | 2868 |
| 2 | M | 275 x 275 | 248.20 | 3282 |
| 3 | C | 275 x 275 | 199.88 | 2643 |
| 3 | G | 275 x 275 | 235.50 | 3114 |
| 3 | M | 275 x 275 | 223.09 | 2950 |
| 4 | C | 300 x 300 | 266.40 | 2960 |
| 4 | G | 300 x 300 | 140.94 | 1566 |
| 4 | M | 300 x 300 | 309.60 | 3440 |

**Supplementary Video 1. Example of a deterrence experiment showing *T. perversa* choice for a control sponge.**

**Supplementary Figure 1. Automatic cell counting method.** (**A**) Starting at the surface, 6 Region Of Interests (ROI 1 - ROI 6, yellow squares) with a depth of 100 μm and a length of 500 μm were selected in each microscopic image. Parameters were adjusted to count spherulous cells (blue outline). (**B**) Samples with longer surface area were collected in 2017. To keep each measured area constant with that in 2016, two regions (Region 1 and Region 2; each region with an area of 600 μm in depth x 500 μm in length) in each microscopic image were selected by avoiding of spongin fibers and aquiferous canals. For each region, the same counting method was applied for 6 ROIs (see A). Then, the average density of spherulous cells from these two regions was calculated for each ROI. (**C**) Comparison between automatic and manual counting method by counting samples from 2016. Both methods showed a similar pattern of the cell density. Manual counting was done by using “Multi-Point” ImageJ tool.

**Supplementary Figure 2. Microscopic section showing shedding and debris in wounded samples collected at 1d.** (**A**) Shedding: microscopic images at 400x showed shedding of spherulous cells (**arrowhead**) out of wounded surface (**s**) and into aquiferous canals (**ca**) in wounded samples (**A1**). TEM-image showing shed spherulous cells (**A2, arrowhead**) out of wounded surface (**s**) and an excreted spherule at the surface (**A2, arrow**). Scale bar= 20 μm (A1), = 5 μm (A2). (**B**) Debris in wounded sponges (100x magnification) (**d**) which were excreted into aquiferous canals (**B1**) or shed out of wound (**B2**). Spongin fibers were also observed to protrude into aquiferous canals or out of the surface (**arrowhead**). Scale bar= 200 μm; **s**= wounded surface; **ca**= aquiferous canal.

**Supplementary Figure 3. Microscopy and photographical recording of recovery after wounding.** (**A**) Example of grazed sample directly after removal of the sea slug (A1). Its osculum (white arrowhead) and the edges between the scar and the intact tissue were regenerated after 3d (A2). (**B**) Example of a microscopic section (400x) of samples from control group (B1) and from grazing group at 1d (B2) and 3d (B3). The surface side was at left of each image. Scales at bottom showed the depth from the surface.

**Supplementary Figure 4. Mass Spectra of unidentified brominated compounds from *A. aerophoba* detected by MALDI-imaging MS.** Five unidentified Br-containing compounds were detected (possibly as $[M+H]^+$ ions) and listed by *m/z* value (from top to bottom panel). The experimental isotopic pattern (left panel), the potential molecular formula (middle panel), and the theoretical isotopic pattern (right panel) for each compound are showed.

**Supplementary Figure 5. MALDI-imaging MS images of ions with an isotopic pattern characteristic of brominated compounds (other than aerophobin-2 and aeroplysinin-1).** Relative abundance, experimental most intense *m/z* in the isotopic pattern, and expected number of Br atoms (in round brackets) are shown for each unidentified brominated compound in the three different treatments (C=control, G=grazing, and M=mechanical damage) for each biological replicate (Replicate; i.e., specimens of the same sponge individual). The relative intensity from 0 to 100 % of each compound is depicted in a colour scale with warmer colours representing relatively higher intensity and colder colours lower intensity of each compound. Scale bar = 1 cm; White-dotted line = broken or cut edges.

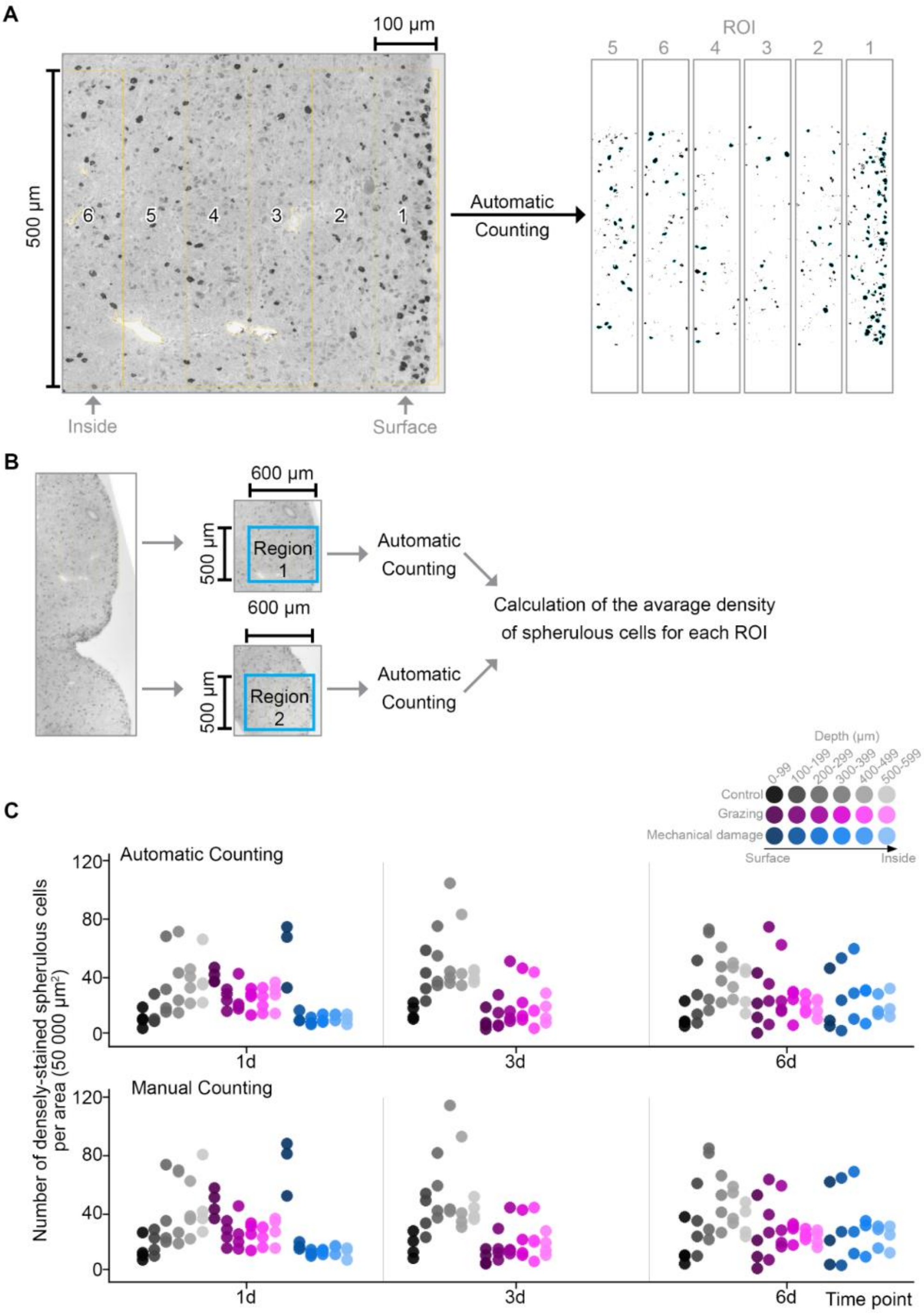


Supplementary Figure 1

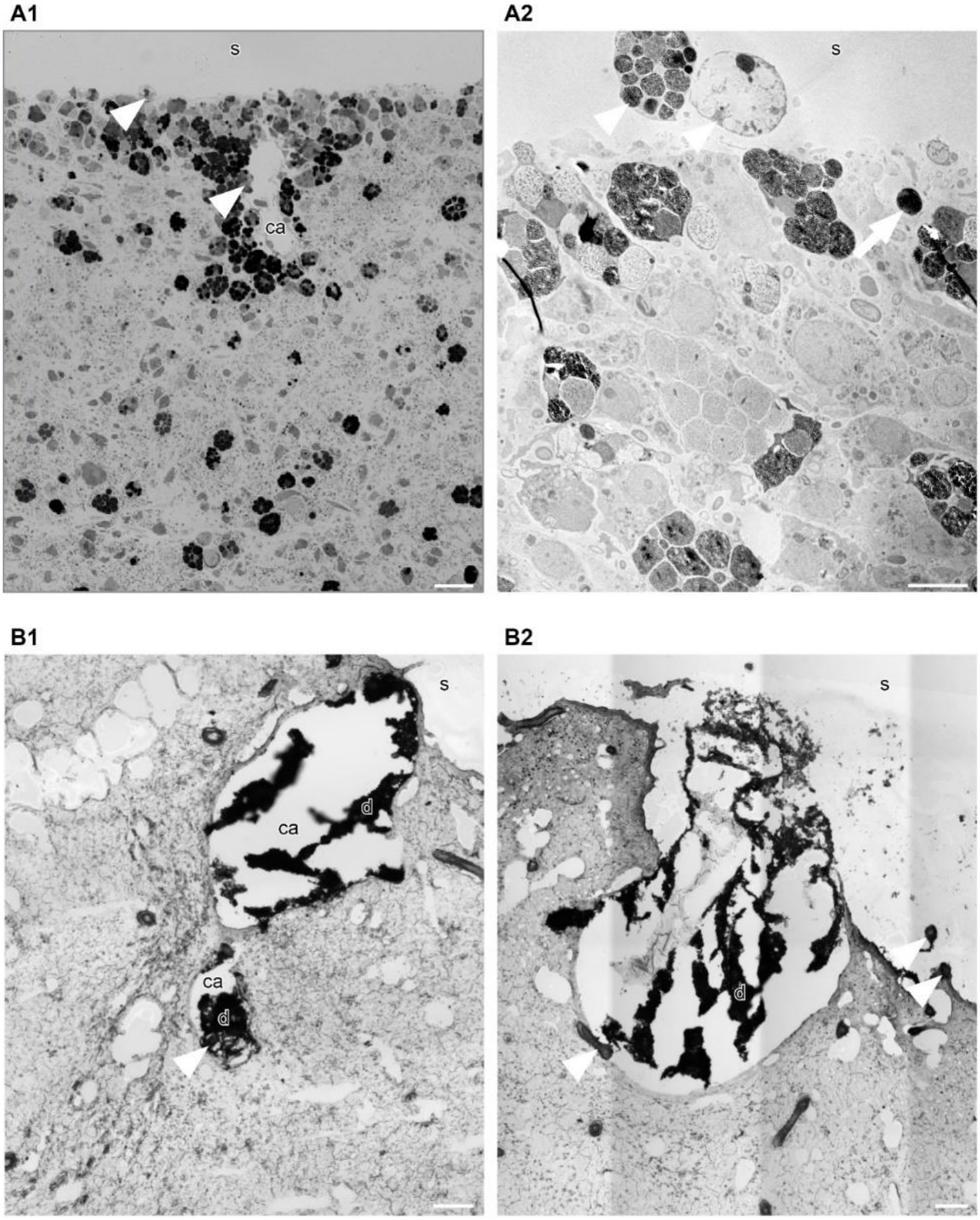


Supplementary Figure 2

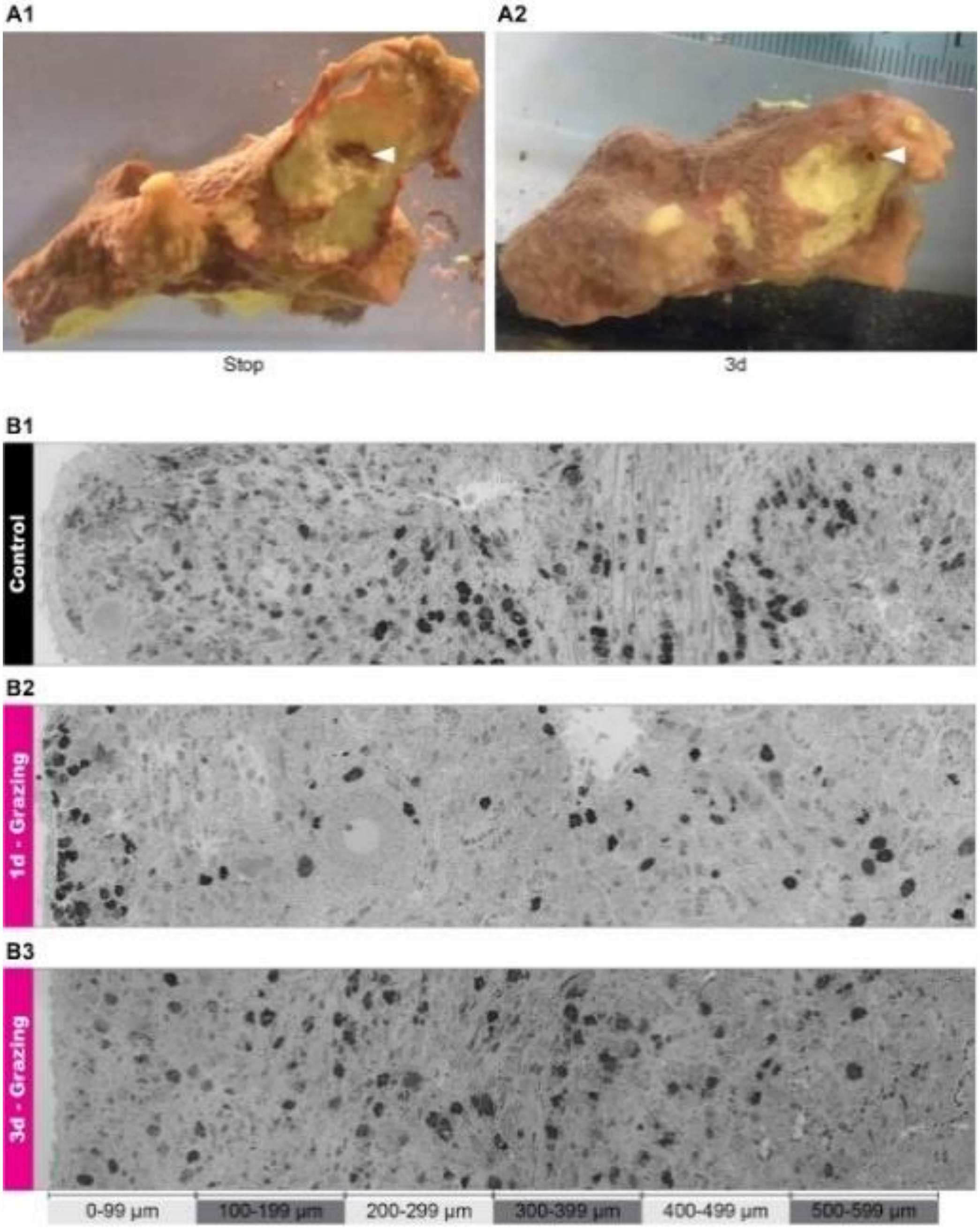


Supplementary Figure 3

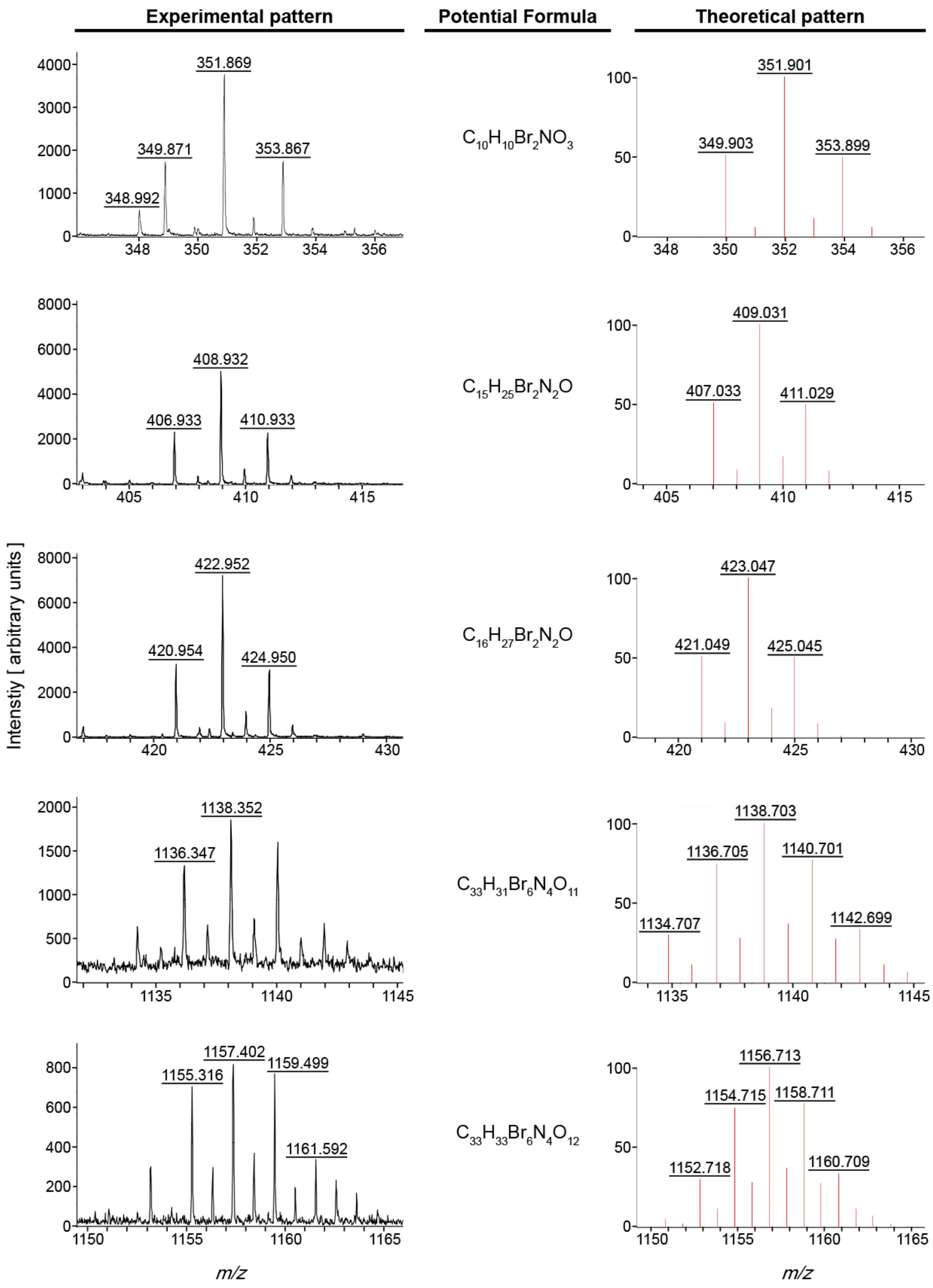


Supplementary Figure 4

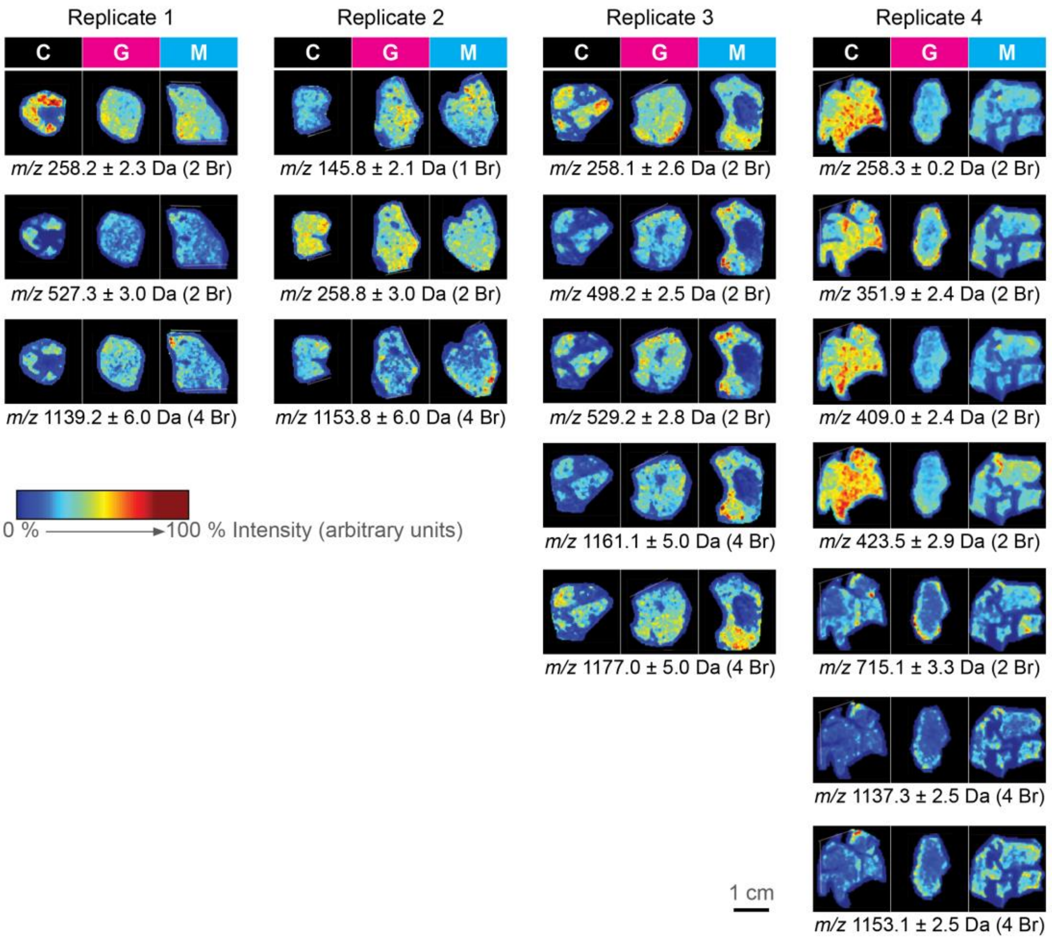


Supplementary Figure 5
